# Programmable multi-kilobase RNA editing using CRISPR-mediated trans-splicing

**DOI:** 10.1101/2023.08.18.553620

**Authors:** Jacob Borrajo, Kamyab Javanmardi, James Griffin, Susan J. St. Martin, David Yao, Kaisle Hill, Paul C. Blainey, Basem Al-Shayeb

## Abstract

Current gene editing approaches in eukaryotic cells are limited to single base edits or small DNA insertions and deletions, and remain encumbered by unintended permanent effects and significant challenges in the delivery of large DNA cargo. Here we describe Splice Editing, a generalizable platform to correct gene transcripts *in situ* by programmable insertion or replacement of large RNA segments. By combining CRISPR-mediated RNA targeting with endogenous cellular RNA-splicing machinery, Splice Editing enables efficient, precise, and programmable large-scale editing of gene targets without DNA cleavage or mutagenesis. RNA sequencing and measurement of spliced protein products confirm that Splice Editing achieves efficient and specific targeted RNA and protein correction. We show that Splice Editors based on novel miniature RNA-targeting CRISPR-Cas systems discovered and characterized in this work can be packaged for effective delivery to human cells and affect different types of edits across multiple targets and cell lines. By editing thousands of bases simultaneously in a single reversible step, Splice Editing could expand the treatable disease population for monogenic diseases with large allelic diversity without the permanent unintended effects of DNA editing.

**One-sentence summary:** CRISPR-guided trans-splicing enables efficient and specific recombination of large RNA molecules in mammalian cells, with broad applications in therapeutic development for genetic diseases and as a research tool for the study of basic RNA biology.

## Introduction

Following transcription, eukaryotic cells extensively reconfigure precursor mRNA (pre-mRNA) to produce the mature mRNA, which constitutes the coding sequence needed by the ribosome to produce protein products. Reactions to process pre-mRNA include 5’ capping, splicing, 3’ end cleavage, and polyadenylation (*1*). The spliceosome, a large ribonucleoprotein (RNP) complex, selectively ligates exons together while excising introns during the splicing process. Splicing occurs *in cis* when the spliceosome ligates an upstream splice donor (SD) to a downstream splice acceptor (SA) within the same pre-mRNA molecule (Figure S1A) (*2*). Most gene products are cis-spliced; however, some gene products naturally rely on RNA trans-splicing (*3*), which occurs when the spliceosome ligates a SD on one molecule to a SA on a different pre-mRNA transcript, creating a single chimeric mRNA molecule (Figure S1B).

Therapeutic trans-splicing, wherein a non-natural trans-splicing reaction is engineered to alter the coding sequence of one or more mRNAs, has the potential to be a widely applicable therapeutic strategy (*4*). Exon exchange via trans-splicing is particularly relevant for monogenic diseases with high allelic diversity since it could, in principle, repair a large spectrum of pathogenic variants in RNA including single nucleotide polymorphisms (SNPs) and large insertions and deletions (indels). As therapeutic changes induced by trans-splicing would be transient and expression of the engineered trans-spliced gene product would remain under endogenous regulation, trans-splicing is predicted to possess a superior safety profile compared to DNA editing and gene therapy (*5, 6*). In contrast, methods using CRISPR-based DNA editing mutagenesis must be carefully designed and evaluated to avoid adverse outcomes. Studies using CRISPR-Cas9 have demonstrated that off-target Cas-based mutations can occur in DNA and RNA using base editors (*7–9*) and in DNA using nucleases (*10, 11*), in addition to chromosomal loss and chromothripsis induced by on-target double-strand breaks (*12, 13*). While attractive as a therapeutic approach, previously reported trans-splicing methods for RNA correction such as SMaRT (Spliceosome-mediated RNA trans-splicing) have relied solely on RNA-RNA hybridization and have suffered from poor editing efficiency and specificity, hindering translation to the clinic (*14*). Here, we describe an RNA editing framework that achieves highly efficient trans-splicing by using RNA-guided proteins to specifically direct a multi-kilobase long repair RNA (repRNA) to the vicinity of the targeted splice junction.

CRISPR-Cas13 is an RNA-guided, RNA-targeting viral defense system that employs two Higher Eukaryotes and Prokaryotes Nucleotide-binding (HEPN) endoRNase domains to cleave the mRNA transcripts of invading viruses within bacteria and archaea. The RNA-targeting ability of CRISPR-Cas13 has been repurposed to develop tools for specific RNA knockdown (*15, 16*), RNA imaging (*17*), targeted RNA base editing (*18, 19*), exon skipping (*20–22*) and targeted RNA methylation (*23*). We envisioned that highly efficient trans-splicing could be achieved using a HEPN-nuclease-deactivated Cas13 variant (dCas13) to recruit a trans-splicing repRNA and simultaneously inhibit cis-splicing by targeting a SA or SD. To accomplish this, we designed specific CRISPR RNA-guided complexes composed of a guide RNA (gRNA), a dCas13 fused to an RNA binding protein (RBP), and a repRNA containing an exon, or group of exons of interest and an engineered intron with RBP-binding hairpins (Figure 1A, B). CRISPR-mediated trans-splicing, which we henceforth refer to as Splice Editing, is achieved by binding the CRISPR-Cas RNP and repRNA complex to a target pre-mRNA. The dCas13-RBP binds to and blocks the cis-SA while simultaneously recruiting the splice repRNA, thus enabling efficient and specific trans-splicing to produce a corrected gene transcript (Figure 1B).

**Figure 1.**
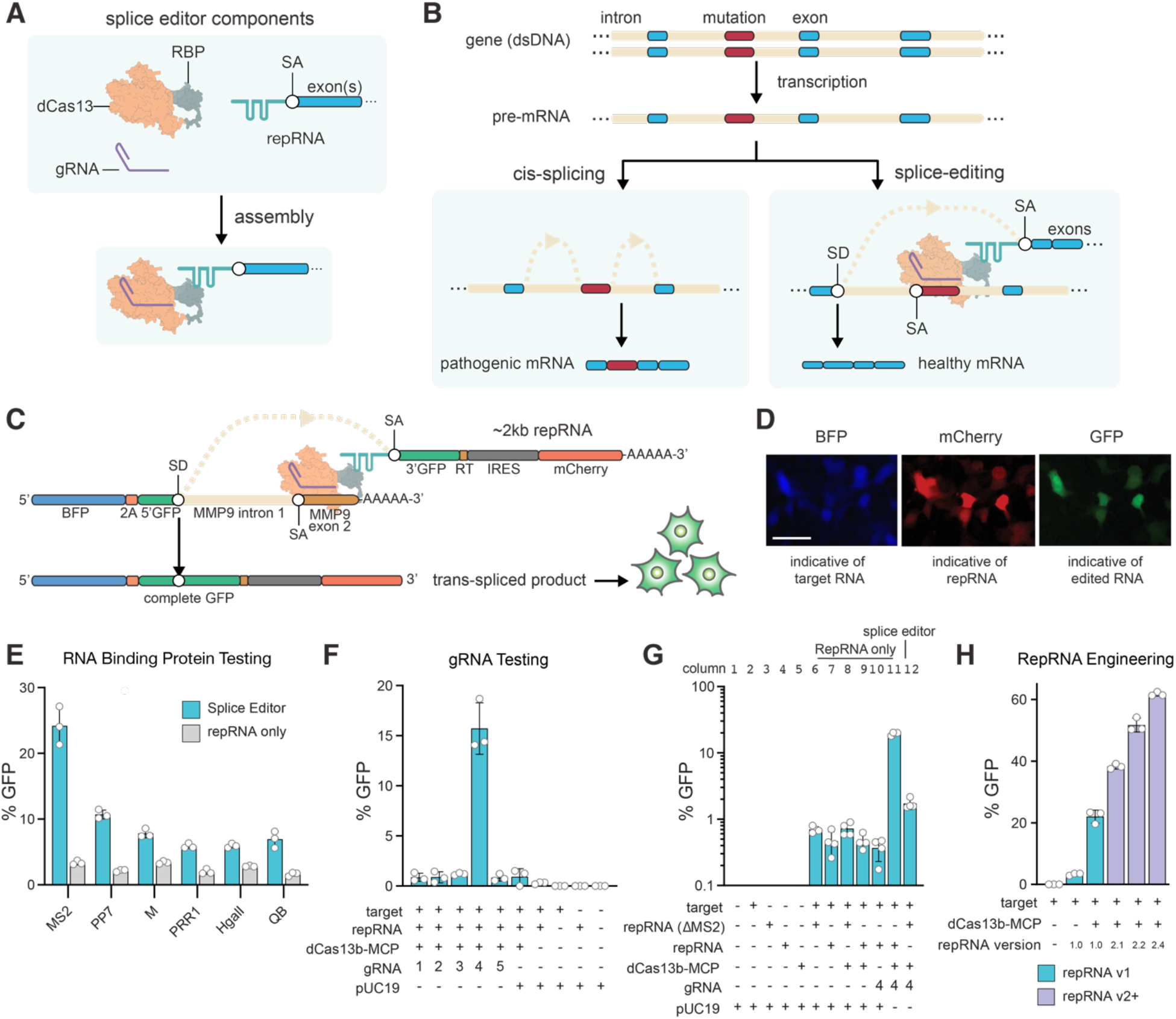
Splice Editing overview, mechanism, and validation. (**A**) Components and assembly of the Splice Editing system. (**B**) Mechanism of RNA cis-splicing and Splice Editing. (**C**) Splice Editing activity in cells is measured by GFP fluorescence, as a complete GFP coding sequence is only present after successful trans-splicing. (**D**) RNA trans-splicing reporter assay, scale bar = 25 μm. (**E**) Activity of targeted RNA trans-splicing using various RBPs in mammalian HEK293FT cells. **(F)** Activity of targeted RNA trans-splicing using various gRNAs with the dCas13b-MCP splice editor in HEK293FT cells (n=3 biological replicates). The full panel is shown in Supplementary Figure S1. **(G)** Examination of GFP editing rate in HEK293FT cells under various conditions at Log scale to resolve differences in trans-splicing activity. Activity in the presence of the Splice Editor was 19.58% ± 1.01% (mean ± SD, n = 4, 2 biological replicates, 2 technical replicates) equating to a 28.03 ± 1.45-fold increase above the repRNA-only condition. Removing the MS2 hairpins (ΔMS2/ MS2-null) from the repRNA (column 12) also showed a small but significant (2.45 ± 0.45-fold (p = 7.3e-4, t-test)) increase in trans-splicing compared to when the dCas13b-MCP and gRNA were not expressed (column 6) **(H)** Effect of engineered repRNAs on Splice Editing activity in mammalian cells. Engineered repRNAs are shown in purple. Increasing interactions between the repRNA and the SE protein components led to >60% editing efficiency.

### Splice Editing enables efficient multi-kilobase RNA editing in mammalian cells

To test Splice Editing in human embryonic kidney 293FT cells (HEK293FT), we constructed a synthetic reporter system wherein a target SD and repair SA RNA (repRNA) must undergo precise trans-splicing to join two GFP exons *in trans* and produce a complete GFP coding sequence (Figure 1C, D). Cells with productive trans-splicing of the reporter can then produce a functional GFP protein product that can be quantified with flow cytometry (Figure S2; See Methods).

We first investigated proof of concept for Splice Editing using the dCas13b variant from *Prevotella sp*. (PspCas13b), fused to a panel of RBPs sourced from RNA bacteriophage genomes. We designed a repRNA containing RBP-binding hairpins, exons encoding GFP and mCherry, and a binding motif (BM) which hybridizes to a site downstream of the SD in Matrix Metallopeptidase 9 (MMP9) intron 1. Using a gRNA targeting the MMP9 splice acceptor site, we observed that MS2 bacteriophage coat protein (MCP) provided the highest level of Splice Editing activity of the RBPs tested, enabling successful insertion of 2 kb at the 3’ end of the target transcript in 24% of cells (Figure 1E, Figure S1C).

### Design rules and engineering for improved Splice Editing

To explore gRNA design rules, we also targeted gRNAs to different regions of the target reporter (Figure S1D) and measured trans-splicing activity. When using a gRNA that targeted the splice acceptor site (gRNA 4), the fraction of cells with Splice Editing-driven GFP signal improved by ∼50-fold (Figure 1F) in comparison to SMaRT, where an RNA motif is solely responsible for hybridizing to the target pre-mRNA to direct trans-splicing. As expected, targeting the CRISPR-Cas system to alternative regions of the target reporter, or excluding dCas13-RBP (Figure S1E, F), did not lead to substantial gains in trans-splicing activity, suggesting that trans-splicing efficiency is improved by blocking the cis-SA. To validate that the observed Splice Editing efficiency was due to the recruitment of the repRNA, we tested a repRNA lacking MS2 hairpins, which led to markedly lower trans-splicing. Interestingly, expressing the MS2-null repRNA along with the other Splice Editing components also showed a smaller but measurable increase in trans-splicing compared to the case where dCas13-RBP and a gRNA were not expressed (Figure 1G). This suggested that binding dCas13 to the splice acceptor alone disfavors cis-splicing, thereby promoting competition by trans-splicing.

To better understand the limiting components in the Splice Editing system, we performed a titration experiment by varying the amount of each component including the target, repRNA, guide RNA, and dCas13-RBP plasmids. Introducing more repRNA led to higher editing efficiency (Figure S3A). However, varying the quantities of guide RNA and dCas13-RBP had little effect over a broad range of concentrations (Figure S3B). Altogether, these results indicated that the repRNA was limiting in our system, as Splice Editing rates were more sensitive to changes in the delivered amount of repRNA. Thus, we posited that changing the number of RBP-binding hairpins in the repRNA could enhance Splice Editing performance by improving the recruitment of the repRNA to the CRISPR-Cas RNP and the target pre-mRNA (Figure S4A). We observed that increasing the number of hairpins led to higher levels of Splice Editing, with repRNA2.4 achieving efficiencies of over 60% (Figure 1H). Further, to explore the impact of protein architecture, we created diverse designs of the protein components, confirming that the most efficient architecture employs a C-terminal RBP fusion with an N-terminal and a linker-embedded Nuclear Localization Signal (Figure S4B).

### Splice Editing is specific to the target pre-mRNA

To characterize the specificity of the Splice Editor, we prepared RNA-seq libraries by specific reverse transcription of the repRNA. Mapped reads showed high specificity (>95%) of the trans-splicing molecules with and without the Splice Editor (Figure 2A), confirming that the binding motif (BM) in the repRNA was specifically addressed to the targeted pre-mRNA. Further, the libraries showed high mapping to the exonic region of the target reporter, with minimal intronic mapping, validating effective trans-splicing (Figure 2B). Full-length RNA-seq on poly-adenylated RNA showed low mapping in the intronic region of the target (MMP9 intron 1) and higher mapping in the exonic regions of the synthetic target (BFP-2A-5’GFP and MMP9 exon 2), confirming expected ongoing cis-splicing of the target (Figure 2C). Deep RNA sequencing showed cis-spliced and trans-spliced exon junctions from the target RNA reporter (Figure S5A). Cis-spliced exon junctions decreased in the presence of the Splice Editor (Figure S5B, columns 11 and 12), further demonstrating that the Splice Editor increases targeted trans-splicing efficiency by attenuating cis-splicing and simultaneously recruiting a repRNA capable of trans-splicing. Furthermore, we did not detect differences in cis-splicing rates in GAPDH, suggesting that the Splice Editing system does not interfere with the cis-splicing of non-targeted pre-mRNAs (Figure S6).

**Figure 2.**
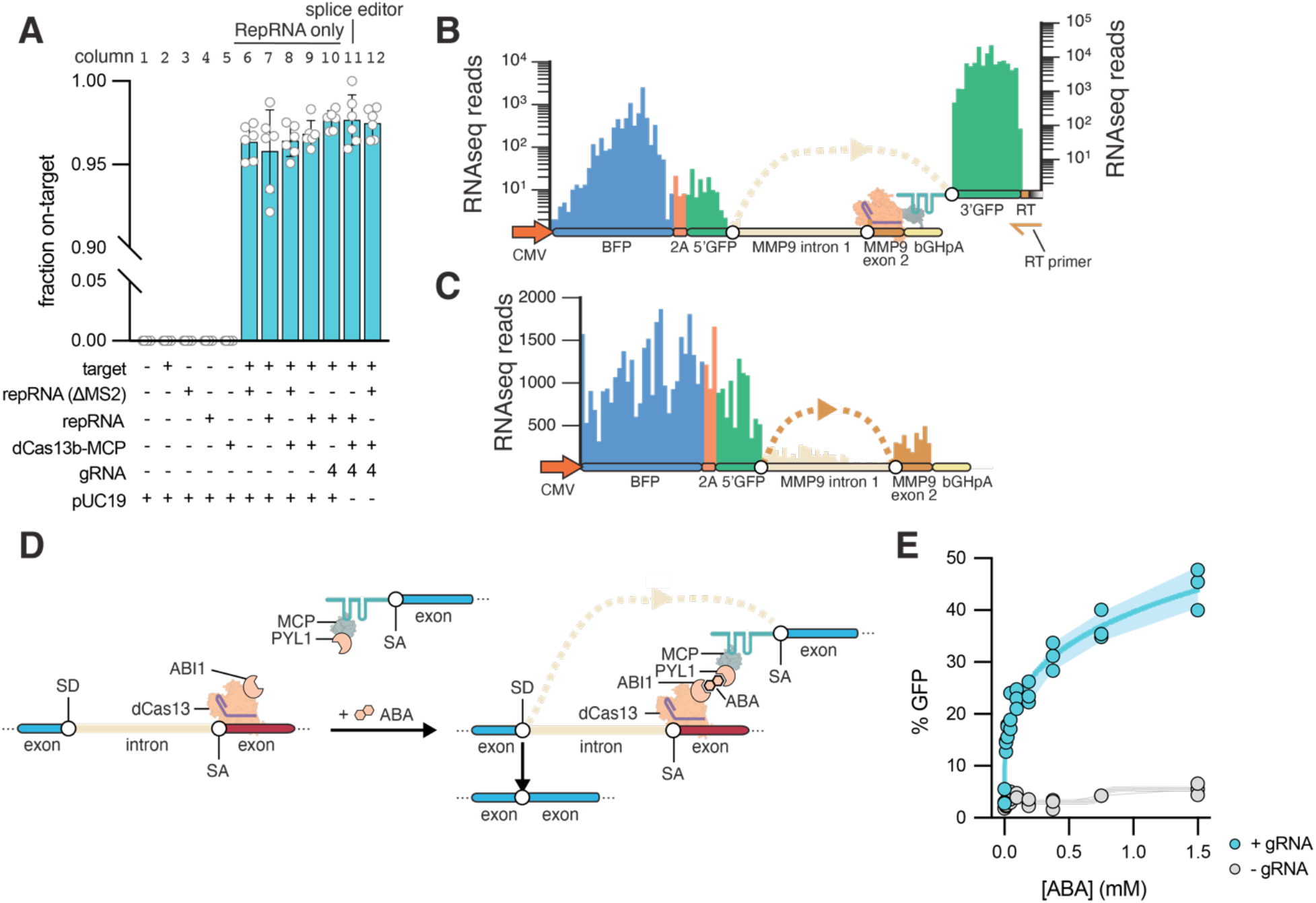
Splice editing characterization. (**A**) Specificity of trans-splicing via RNA-seq. (**B**) RNA-seq prepared from the reverse transcription site (RT) demonstrates trans-splicing. (**C**) Validation of reporter cis-splicing via RNA-seq. (**D**) Chemically inducible Splice Editing system using abscisic acid (ABA). (**E**) Splice editor activity is tunable by abscisic acid dosage when a guide RNA is present and remains at background levels in the absence of a guide RNA. Error bands indicate Standard Deviation.

### Tunable and chemically-induced Splice Editing

Next, we explored the possibility of finely tuning Splice Editor activity with an exogenous molecule, with the rationale that RNA correction levels could be modulated by adjusting the dosage of an administered small molecule. To model control by a small molecule, we developed a chemically-induced Splice Editor using the ABI1/PYL1 system, which undergoes heterodimerization when binding to the plant phytohormone abscisic acid (ABA) (*24*) (Figure 2D). We fused Cas13b to ABI1, and fused MCP to PYL1, both via glycine-serine linkers within a single 2A-linked expression cassette. Using our reporter system, we observed fine control of Splice Editing efficiency upon ABA titration after cell transfection (Figure 2E) with >40% of cells transitioning from GFP- to GFP+ at 1.5 mM ABA. Thus, we find that small molecule control can support tunable and reversible trans-splicing, and further demonstrate that the activity of the system is largely dependent on the proximity of the repRNA relative to the target pre-mRNA.

### CRISPR-mediated internal exon replacement

The ability to rewrite internal exons is a desirable feature for various RNA editing contexts. Having demonstrated the successful replacement of the 3’ end of mRNAs, we sought to determine whether Splice Editing is capable of internal exon replacement. We designed a reporter system to monitor trans-splicing events that replace an internal exon. For this, we split GFP into three exons, utilizing GFP exons 1 and 3 in the target RNA molecule design. To simulate a pathogenic exon, we placed MMP9 exon 2 between GFP exons 1 and 3 on the target molecule, along with corresponding flanking introns: MMP9 intron 1 and MMP9 intron 2. The internal repRNA was designed to have GFP exon 2 flanked by two synthetic introns, each with an MS2 hairpin (Figure S7A). We monitored the expression of the target transcript by expression of BFP. Upon internal exon repair, GFP exon 2 from the repRNA is trans-spliced to create a complete GFP mRNA (Figure S7B). With the internal trans-splicing reporter system, we observed gRNA-dependent internal exon replacement with no detectable exon replacement without the complete Splice Editor (Figure S7C). These results demonstrate that Splice Editing is capable of internal exon replacement, which employs both 3’ and 5’ trans-splicing.

### Phylogenetic and structural insights of novel miniature CRISPR-Cas effectors

Despite the validation of Splice Editing as efficient and specific in mammalian cells, the therapeutic delivery of the Cas13-based fusion proteins presented a challenge, since the sizes of reported Cas13 genes as fusions exceed the packaging capacity of most adeno-associated virus (AAV) serotypes (4.7kb). To address Splice Editor delivery limitations and enable *in vivo* multi-kb nucleic acid editing, we sought to discover and characterize the diversity of miniature Cas13 proteins that may be packaged as a splice-editing fusion protein within an AAV delivery vector. Using metagenomic analysis of microbial samples, we encountered new variants of Cas13x (Cas13bt) (*25, 26*) but also identified a phylogenetically distinct Cas13 protein family ∼40-50% smaller than LbaCas13a (*27*) and ∼15-20% smaller than EsiCas13d (*28*), which we dubbed Cas13e (Figure 3A). These CRISPR-Cas systems were primarily encoded in genomes of new sulfur-oxidizing *Epsilonproteobacteria* species (Figure 3A), where their CRISPR arrays were found to target transcripts of bacteriophages and plasmids found within the same samples (Figure 3B, Supplementary Table S1).

**Figure 3.**
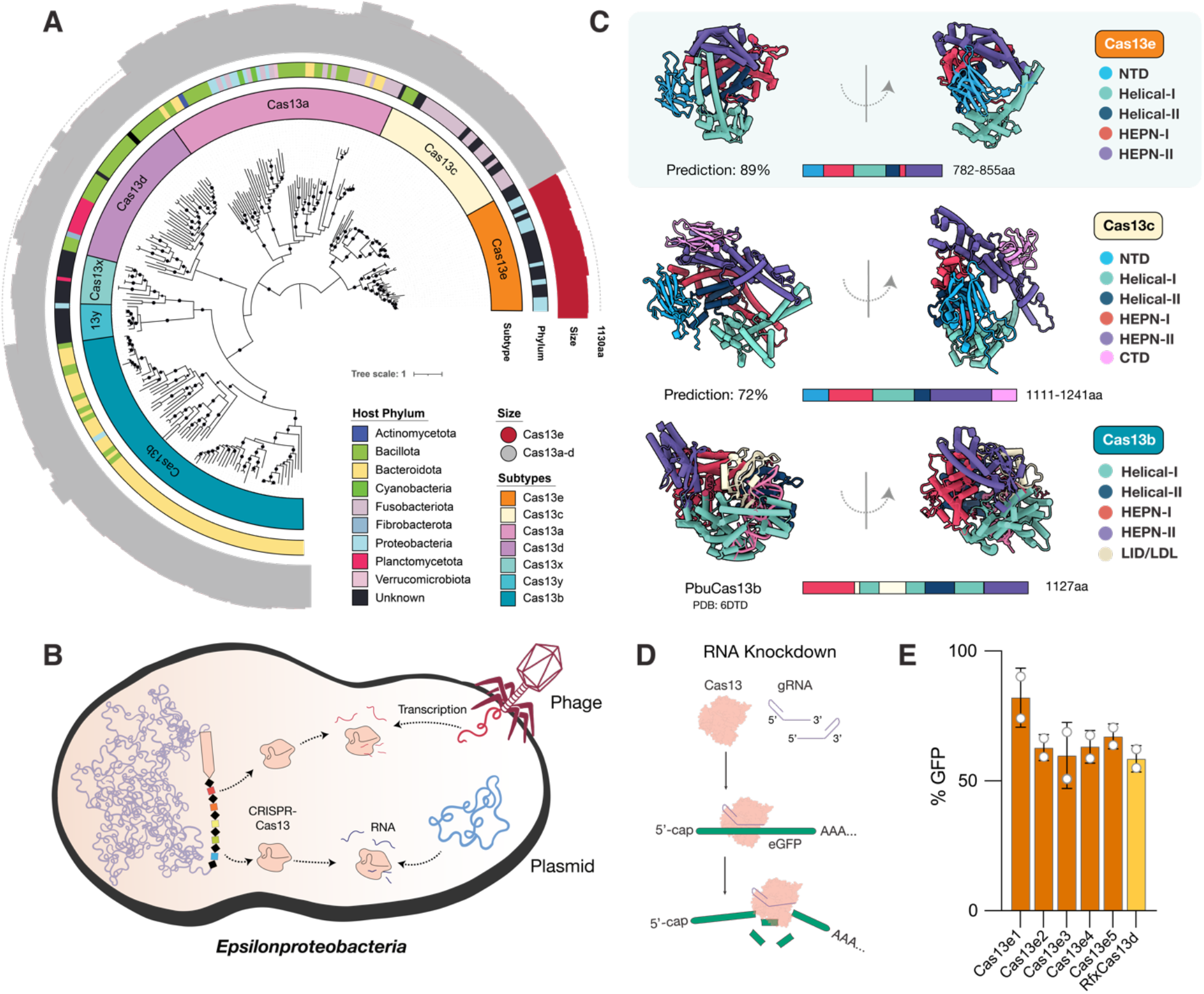
**(A)** Maximum Likelihood phylogenetic reconstruction of previously reported and newly discovered CRISPR-Cas13 systems. The inner to outer rings display the CRISPR–Cas subtype, bacterial phylum in which specific protein representatives were found, and protein size with a dotted circle outlining the average Cas13 size. Bootstrap values >90 are signified by black dots. The phylogenetic distance of the new Cas13 clade from Cas13c is comparable to the distance between Cas12c and Cas12d families. Given the miniaturized size, evolutionary and structural divergence from previously described systems, as well as the difference in functional activity in cellular contexts, we refer to the novel group described herein as Cas13e. **(B)** Cell diagram of bacterium–phage interactions that involve CRISPR-Cas13e targeting. Arrows indicate CRISPR– Cas targeting of the phage and plasmid genomes identified from metagenomes of the same samples or the same site. **(C)** ColabFold-based predicted protein structures and domain organizations of Cas13e and Cas13c representatives relative to EsiCas13d (PDB:6E9F) **(D)** Schematic of RNA knockdown strategy to validate the activity of Cas13e in mammalian cells **(E)** Relative GFP fluorescence (MFI targeting crRNA/MFI non-targeting crRNA) of HEK293T-GFP cells transfected with plasmids expressing Cas13e or RfxCas13d and GFP-targeting crRNA, measured by flow cytometry. Percent GFP detected in mammalian cells relative to non-targeting negative controls.

The most closely related enzyme to the Cas13e nuclease is Cas13c (15% identity; Figure 3A), which remains the least characterized Cas13 and the rarest to date, with few homologs reported. Little is known about the Cas13c enzyme in terms of structure or mechanism, and it has not been successfully employed for RNA editing in mammalian cells (*18, 25*). To compare the domain architecture of the two Cas families and infer how they differ from other Cas13 proteins, we produced high-confidence predictions of their structures using ColabFold (Figure 3C, Figure S8). As has been reported for other single-effector (Class 2) Cas proteins, the Cas13e predicted structure is bi-lobed. Cas13e consists of two HEPN domains in the Nuclease (NUC) lobe for cleavage of RNA targets, and a Recognition (REC) lobe consisting of α-helical domains and an N-terminal domain (NTD) that may be used to bind the CRISPR repeat of the gRNA. As opposed to the NTD composed of α helices in Cas13a, or short α helices flanking a β sandwich region in Cas13d, the NTD in Cas13e and Cas13c appears to be composed almost entirely of β sheets. It is expected that bound crRNA and target RNA in the Cas13e RNP would be coordinated within the large, positively charged channel through the center of the enzyme (Figure 3C; Figure S8). Interestingly, we discovered the presence of a novel C-terminal domain (CTD) in the NUC lobe of Cas13c that is not found in previously reported Cas13 systems and is composed primarily of β sheets. In addition to the miniaturization of the NTD, HEPN, and helical domains in Cas13e compared to Cas13c, there was no homologous domain structure to the CTD in Cas13e. Overall, phylogenetic and structural insights support the distinctiveness of the miniature Cas13e effector as compared to previously described Cas proteins.

### Miniature Splice Editors using CRISPR-Cas13e

Given the relatively small size of Cas13e and its sequence divergence from Cas13 systems successfully employed in mammalian cells (<7%), we first sought to validate its activity in HEK293 cells. We tested the ability of the native Cas13e enzymes to target and cleave GFP transcripts as compared to RfxCas13d using different guide orientations (Figure 3D). Cas13e3 exhibited the most robust RNA knockdown activity, with a gRNA orientation that was the reverse of that of Cas13x (Cas13bt) (Figure 3E; Figure S8). We thus created nuclease-inactive Cas13e3 with R244A/H249A and R704A/H709A mutations in the HEPN1 and HEPN2 domains, respectively. Using dCas13e3, we were able to create a Splice Editor (SE2) that is 22% smaller than SE1, which enables the packaging of the full fusion protein into AAVs for delivery to mammalian cells. We first demonstrated the efficacy of the SE2 system by targeting the MMP9 splice acceptor site, leading to multi-kilobase 3’ replacement of RNA in HEK293 and HepG2 cells, demonstrating that Splice Editing is active in a variety of human-derived cell lines (Figure 4C).

**Figure 4.**
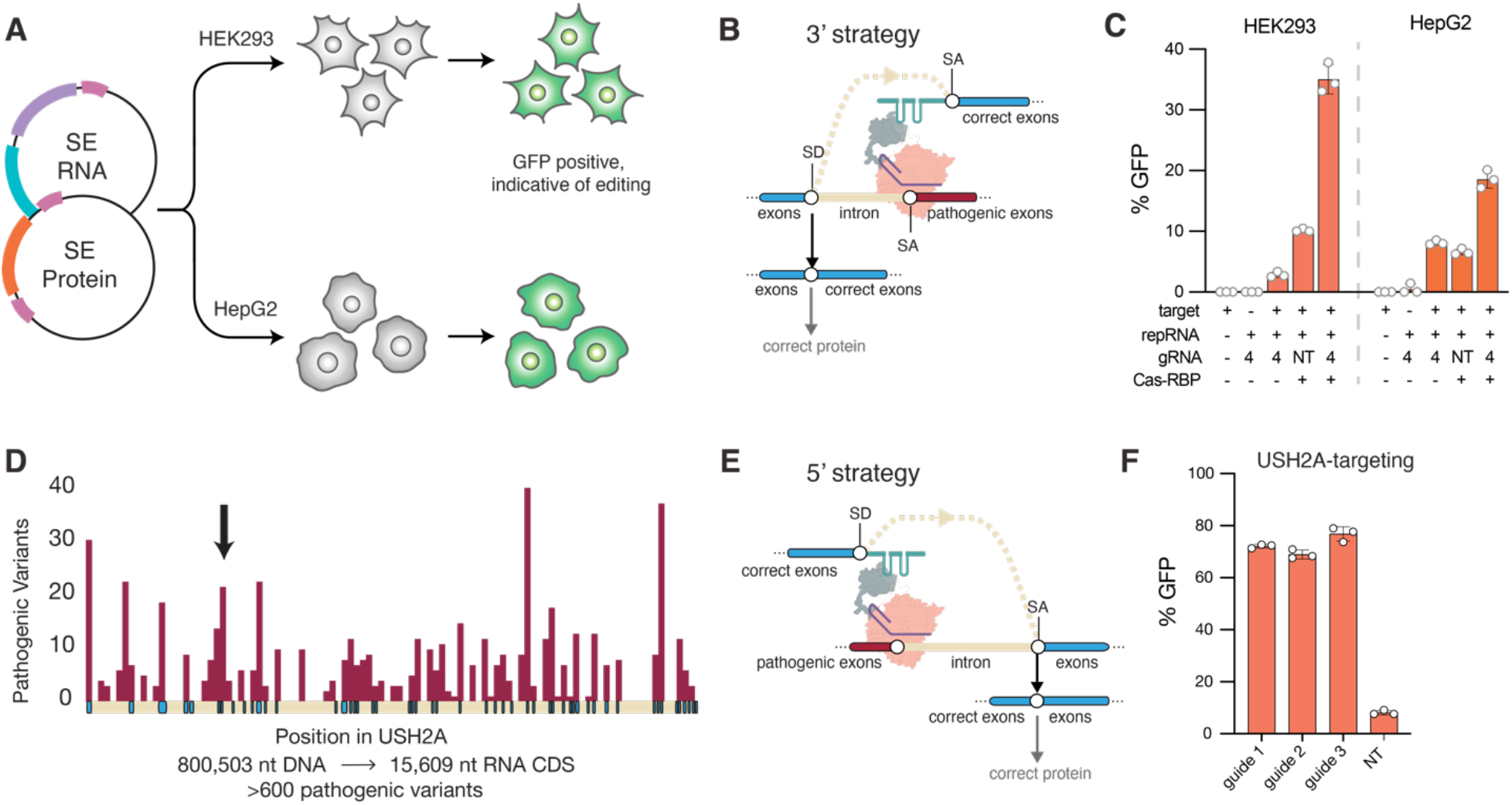
**(A)** Schematic of experimental workflow for evaluating the performance of SE2 for RNA replacement as plasmids in different mammalian cell types. **(B)** The strategy for RNA replacement at the 3’ end. **(C)** 3’ editing performance as applied to a MMP9 reporter using SE2 constructs with targeting and non-targeting (NT) guides in HEK293 and HepG2 cells. **(D)** The number of pathogenic mutations is plotted in maroon per position in the USH2A gene, with exons shown in blue and introns shown in light cream color. Over 600 pathogenic variants occur throughout the full length transcript. The position of exon 13 is signified by a black arrow. **(E)** A proposed strategy for the correction of the 5’ end of a target RNA. **(F)** 5’ editing as applied to a USH2A reporter, with gRNAs targeting intron 13, vs. a non-targeting guide where activity is driven by the presence of a repair RNA without a directed binding motif.

### Efficient and programmable 5’ RNA replacement in alternative gene targets

Splice Editing is particularly well-suited for monogenic diseases where the affected gene is beyond the packaging limit of AAV and possesses high allelic diversity in the patient population, and a single Splice Edit can address a broad collection of disease alleles. Of these, the USH2A gene has been recognized as one of the most challenging targets for gene therapy (*29*). Mutations in the human USH2A gene account for most cases of Usher Syndrome type II, a heterogeneous autosomal recessive disorder characterized by progressive retinitis pigmentosa and sensorineural hearing deficiencies, resulting in deaf-blindness in patients (*30*). Development of genetic therapies via AAV-mediated supplementation has been hindered due to the large size of the USH2A gene (800 kb) and open reading frame (15.6 kb) compared to the AAV packaging capacity (4.7 kb) (Figure 4D). Furthermore, pathogenic variation in USH2A is too heterogeneous to be compatible with existing gene-editing approaches, with >600 genetic variants spanning missense, nonsense, insertion, deletion, and frame-shift mutations throughout its 72 exons (*29, 31*) (Figure 4D). Using the minimal Cas13e-based SE2, we aimed to validate Splice Editing as a corrective approach for USH2A transcripts and demonstrate the possibility of replacement of the 5’ end of an RNA target (Figure 4G). To this end, we designed gRNAs to bind intron 13 within an USH2A reporter, and a repRNA that would enable GFP fluorescence upon transcript correction with 5’ replacement. Via transient transfection into HEK293 cells, we observed Splice Editing efficiencies up to ∼80% (Figure 4F).

## Discussion

Splice Editing represents the first CRISPR-guided platform that enables multi-kilobase correction of transcripts. We have demonstrated that this approach is functional in several mammalian cell types, achieving >97% specificity to the gene transcripts of interest in the process. This offers a strong alternative strategy to CRISPR-Cas9 or DNA-based approaches for sequence replacement that have suffered from substantial and permanent off-target editing (*7–11*) as well as unintended permanent effects of on-target editing (*12, 13*). Furthermore, the lack of a Protospacer Adjacent Motif (PAM) requirement for successful Cas13 targeting dramatically increases the potential target sites within a given sequence relative to a DNA editor.

One key advantage is that Splice Editing does not rely on endogenous DNA repair pathways to generate desired editing outcomes. Instead, it co-opts the spliceosome, which is operational in all mammalian cells with active transcription. It can thus offer particular editing advantages in post-mitotic cells such as neurons, cardiomyocytes, and retinal cells. In such contexts, DNA editing approaches have been relatively inefficient due to reliance on pathways with low activity such as base-excision repair, mismatch repair, or homologous recombination (*32, 33*).

Many diseases are caused by mutations in genes whose size exceeds the packaging limit of AAV, and thus delivery of a full-length therapeutic protein by a single or dual AAV vector (gene therapy) is not a viable strategy. By replacing large segments of the RNA, Splice Editing can provide an especially advantageous strategy for genetic correction in such cases. Moreover, as RNA is a transient molecule, the editing effect is spontaneously reversed by cessation of Splice Editor activity, a feature that alleviates multiple concerns observed in DNA editing, DNA recombination, or gene therapy approaches. For instance, once gene therapy has been delivered into a target tissue, it is difficult to regulate the levels of expression. Tight and appropriate regulation of healthy gene expression is essential for alleviating many diseases; too much expression may have toxic effects, and too little expression may not impart the intended benefits of the therapy (*5, 6*). In contrast, Splice Editing directly rewrites large segments of an already highly and homeostatically regulated gene product. Such endogenous regulatory control improves safety by constraining the expression of trans-products near the natural range and may improve efficacy by retaining appropriate dynamic expression control in response to local signals. We further demonstrated that this approach could be modulated using an exogenous small molecule.

In this framework, a single Splice Editor configuration is sufficient to enable programmable and efficient therapeutic correction of numerous mutations at once via the replacement of one or more exons, which provides significant benefits over ASO, shRNA, and several CRISPR-based DNA editing technologies. Such competing methods target single mutations at a time and would require individual therapies for each disease subpopulation (*34*) with the inequitable likelihood that patients with rare alleles are left behind. Here, we show that Splice Editing enables facile programmable insertion or replacement of 3’, 5’, or internal exons to target any region of an RNA molecule. This provides a consolidated path for therapeutic development addressing large sets of common and rare pathogenic mutations (*35*). Here, we’ve described compact CRISPR-Cas13 systems that are active as Splice Editors and are compatible with AAV-based delivery for *in vivo* applications. Programmable insertion of thousands of bases of RNA into endogenous transcripts in a single step represents a promising approach that could address most loss-of-function mutations across 5000 genetic diseases in a safe and reversible manner.

## Methods

### Cell culture

HEK293FT (Thermo Fisher Scientific Cat. No. R70007) and HepG2 (ATCC Cat. No. HB-8065) were cultured in a humidified incubator at 5% CO2 at 37 C using high-glucose DMEM (Invitrogen) supplemented with 10% FBS, 1% penicillin/streptomycin, and 1% GlutaMAX. Cell cultures were kept at low passage (<20) and regularly tested for mycoplasma contamination.

### Cloning

Plasmids were synthesized by Twist or Vector Builder, and genes were synthesized by Twist or IDT. The RBP sequences were synthesized by IDT and inserted with Golden Gate assembly. gRNAs were ordered as oligos from IDT and were hybridized and phosphorylated for Golden Gate assembly.

### Transfections

HEK293FT cells were transfected in a 12-well plate format, with a total of 1250 ng of DNA and 4 μL of Lipofectamine 2000 per condition. Each construct was one-fourth (312.5 ng) of the total DNA transfection, with the exception of pUC19, which was used as non-coding dummy DNA in conditions where fewer than four components were delivered, such that there would be a total of 1250 ng of DNA in each transfection. Media was changed 6 hours after the transfection, and the cells were analyzed via flow cytometry (Beckman Coulter CytoFLEX platform) 48 hours after the transfection.

### Internal exon transfections

HEK293FT cells were transfected using Lipofectamine 2000 per the manufacturer’s instructions and cells were transfected in a 96-well format (100 ng per well) with four biological replicates for each condition. Flow cytometry was conducted 48 hours after transfection. Several gRNAs targeting the splice donor (SD) were designed (SD targeting gRNAs 1-5), and delivered in conjunction with a splice acceptor (SA) targeting gRNA. In the absence of gRNAs, no GFP+ cells were detected. However, upon introduction of gRNAs, GFP+ cells were detected and the fraction of GFP+ cells varied depending on the placement of the gRNA relative to the splice donor.

### Abscisic acid (ABA) inducible Splice Editing

HEK293FT cells were transfected using Lipofectamine 2000 per manufacturer’s instructions in 96-well format. 6 hours after transfection, the media was changed and replenished with media containing abscisic acid (Sigma cat. no. A1049) and was titrated across multiple samples with 3 biological replicates per ABA concentration. 72 hours after transfection, the cells were measured via flow cytometry (Beckman Coulter CytoFLEX platform).

### RNA knockdown experiments in mammalian cells

For each Cas ORF, an SV40 NLS sequence was added to the N-terminus (MSPKKKRKVEAS), and an SV40 NLS with an appended HA tag was added to the C-terminus (GSGPKKKRKVAAAYPYDVPDYA). The resulting ORFs were then mammalian codon-optimized and synthesized into mammalian expression vectors. Transcription was driven by a CMV promoter (CGCCCCATTGACGCAAATGGGCGGTAGGCGTGTACGGTGGGAGGTCTATATAAGCA GAGCTGGTTTAGTGAACCGTCAGATC) and an SV40 polyadenylation signal was used to terminate transcription (CAGCTCACAGACATGATAAGATACATTGATGAGTTTGGACAAACCACAACTAGAATGCAGTGAAAAAAATGCTTTATTTGTGAAATTTGTGATGCTATTGCTTTATTTGTAACCATTATAAGCTGCAATAAACAAGTTAACAACAACAATTGCATTCATTTTATGTTTCAGGTTCAGGGGGAGGTGTGGGAGGTTTTTTAAAGCAAGTAAAACCTCTACAAATGTGGTATTGGC). A plasmid expressing eGFP and plasmids expressing Cas13e proteins and gRNAs were co-transfected into HEK293T cells (bio reps n=2) using Lipofectamine 2000 per manufacturer’s instructions. Protein expression was confirmed by observing fluorescence via fluorescent microscopy on a Bio-rad ZOE Fluorescent Cell Imager, and flow cytometry was performed 48 hours following transfection on a Sartorius iQue3. Cell and singlet gating was performed with Sartorius iQue3 instrument software, and the median GFP intensity of singlets was calculated in Python. Results were plotted using Matplotlib (https://matplotlib.org/). gRNAs were designed for Cas13e systems to target multiple sites across the coding sequence of eGFP in HEK293T cells, to be tested alongside a scrambled non-targeting guide, demonstrating re-programmability and specificity of the RNase activity of these CRISPR-Cas systems towards different targets. Several gRNA orientations were tested to evaluate the correct structure of the CRISPR RNA of Cas13e systems. gRNA structure for Cas13e systems was determined to consist of a CRISPR repeat at the 5’ end of the RNA followed by a spacer sequence complementary to the target RNA. In contrast, the gRNA structure for Cas13e systems was determined to consist of a spacer sequence complementary to the target RNA on the 5’ end of the gRNA, followed by a CRISPR repeat at the 3’ end.

### Metagenome-assembled genome construction, binning, and phylogenetic classification of Cas13 genomes

Briefly, samples were assembled with metaSPAdes v3.15.4 (*37*). Proteins were identified using Prodigal v2.6.3 (*38*) and aligned to custom Cas13 profiles. Reference proteins were aligned with Cas13 hits using MAFFT (*39*), and phylogenetic trees were built using IQTree (*40*). Samples containing one or more significant matches to Cas13 systems were binned into metagenome-assembled genomes (MAGs) using metabat2 v2.16 (*41*), MaxBin v2.2.7 (*42*), and CONCOCT v1.1 (*43*) in parallel. Completeness and contamination were estimated using CheckM v1.2.2 (*44*) using lineage-specific markers. Bins recovered from all three tools for each sample were refined and merged using DAS Tool v1.1.6 (*45*). Medium and high-quality MAGs containing Cas13e (≥60% completeness and ≤10% contamination based on essential genes) from all samples were taxonomically classified using GTDB-tk using the “classify_wf” setting. In addition, bin phylogeny was determined by placing these genomes into a phylogenetic tree of existing *Epsilonproteobacteria* genomes built from a concatenated set of rRNA genes. CRISPR spacers from Cas13 genomes were searched against metagenomes using BLAST and hits with ≥20nt of matches and ≤4 mismatches were retained for analysis.

## Author contributions

This study was conceived and co-led by J.D.B. and B.A-S. Experiments were performed by J.D.B., K.J., S.S., D.Y., K.H. with input and supervision from B.A-S. B.A-S and J.G. performed bioinformatic discovery and analyses. B.A-S and J.D.B wrote the manuscript with input from K.J. and P.C.B.

## Acknowledgments

The authors thank Dr. Gene Yeo, Dr. Melissa Jurica, Dr. David Schaffer, and Dr. Fei Chen for comments on the study, and Dr. David Colognori for comments on the manuscript. P.C.B was supported by Burroughs Wellcome Fund CASI Award and the NIH Director’s New Innovator Award (DP2HL141005). J.D.B. was supported in part by the HHMI Gilliam Fellowship, the NSF Graduate Research Fellowship, and the Viterbi Fellowship.

## References

1. D. L. Bentley, Coupling mRNA processing with transcription in time and space. Nat. Rev. Genet. 15, 163–175 (2014).

2. M. M. Scotti, M. S. Swanson, RNA mis-splicing in disease. Nat. Rev. Genet. 17, 19–32 (2016).

3. G. Flouriot, H. Brand, B. Seraphin, F. Gannon, Natural Trans-spliced mRNAs Are Generated from the Human Estrogen Receptor-α (hERα) Gene *. J. Biol. Chem. 277, 26244–26251 (2002).

4. R. G. Pergolizzi, R. G. Crystal, Genetic medicine at the RNA level: modifications of the genetic repertoire for therapeutic purposes by pre-mRNA trans-splicing. C. R. Biol. 327, 695–709 (2004).

5. R. M. Payne, Gene therapy for Friedreich ataxia: Too much, too little, or just right? Mol Ther Methods Clin Dev. 25, 1–2 (2022).

6. A. M. Monteys, A. A. Hundley, P. T. Ranum, L. Tecedor, A. Muehlmatt, E. Lim, D. Lukashev, R. Sivasankaran, B. L. Davidson, Regulated control of gene therapies by drug-induced splicing. Nature. 596, 291–295 (2021).

7. E. McGrath, H. Shin, L. Zhang, J.-N. Phue, W. W. Wu, R.-F. Shen, Y.-Y. Jang, J. Revollo, Z. Ye, Targeting specificity of APOBEC-based cytosine base editor in human iPSCs determined by whole genome sequencing. Nat. Commun. 10, 5353 (2019).

8. E. Zuo, Y. Sun, W. Wei, T. Yuan, W. Ying, H. Sun, L. Yuan, L. M. Steinmetz, Y. Li, H. Yang, Cytosine base editor generates substantial off-target single-nucleotide variants in mouse embryos. Science. 364, 289–292 (2019).

9. J. Grünewald, R. Zhou, S. P. Garcia, S. Iyer, C. A. Lareau, M. J. Aryee, J. K. Joung, Transcriptome-wide off-target RNA editing induced by CRISPR-guided DNA base editors. Nature. 569, 433–437 (2019).

10. Y. Fu, J. A. Foden, C. Khayter, M. L. Maeder, D. Reyon, J. K. Joung, J. D. Sander, High-frequency off-target mutagenesis induced by CRISPR-Cas nucleases in human cells. Nat. Biotechnol. 31, 822– 826 (2013).

11. Y. Lin, T. J. Cradick, M. T. Brown, H. Deshmukh, P. Ranjan, N. Sarode, B. M. Wile, P. M. Vertino, F. J. Stewart, G. Bao, CRISPR/Cas9 systems have off-target activity with insertions or deletions between target DNA and guide RNA sequences. Nucleic Acids Res. 42, 7473–7485 (2014).

12. S. Papathanasiou, S. Markoulaki, L. J. Blaine, M. L. Leibowitz, C.-Z. Zhang, R. Jaenisch, D. Pellman, Whole chromosome loss and genomic instability in mouse embryos after CRISPR-Cas9 genome editing. Nat. Commun. 12, 5855 (2021).

13. M. L. Leibowitz, S. Papathanasiou, P. A. Doerfler, L. J. Blaine, L. Sun, Y. Yao, C.-Z. Zhang, M. J. Weiss, D. Pellman, Chromothripsis as an on-target consequence of CRISPR-Cas9 genome editing. Nat. Genet. 53, 895–905 (2021).

14. A. Berger, S. Maire, M.-C. Gaillard, J.-A. Sahel, P. Hantraye, A.-P. Bemelmans, mRNA transsplicing in gene therapy for genetic diseases. Wiley Interdiscip. Rev. RNA. 7, 487–498 (2016).

15. O. O. Abudayyeh, J. S. Gootenberg, P. Essletzbichler, S. Han, J. Joung, J. J. Belanto, V. Verdine, D. B. T. Cox, M. J. Kellner, A. Regev, E. S. Lander, D. F. Voytas, A. Y. Ting, F. Zhang, RNA targeting with CRISPR–Cas13. Nature. 550, 280–284 (2017).

16. H.-H. Wessels, A. Méndez-Mancilla, X. Guo, M. Legut, Z. Daniloski, N. E. Sanjana, Massively parallel Cas13 screens reveal principles for guide RNA design. Nat. Biotechnol. 38, 722–727 (2020).

17. L.-Z. Yang, Y. Wang, S.-Q. Li, R.-W. Yao, P.-F. Luan, H. Wu, G. G. Carmichael, L.-L. Chen, Dynamic Imaging of RNA in Living Cells by CRISPR-Cas13 Systems. Mol. Cell. 76, 981–997.e7 (2019).

18. D. B. T. Cox, J. S. Gootenberg, O. O. Abudayyeh, B. Franklin, M. J. Kellner, J. Joung, F. Zhang, RNA editing with CRISPR-Cas13. Science. 358, 1019–1027 (2017).

19. O. O. Abudayyeh, J. S. Gootenberg, B. Franklin, J. Koob, M. J. Kellner, A. Ladha, J. Joung, P. Kirchgatterer, D. B. T. Cox, F. Zhang, A cytosine deaminase for programmable single-base RNA editing. Science. 365, 382–386 (2019).

20. S. Konermann, P. Lotfy, N. J. Brideau, J. Oki, M. N. Shokhirev, P. D. Hsu, Transcriptome Engineering with RNA-Targeting Type VI-D CRISPR Effectors. Cell. 173, 665–676.e14 (2018).

21. M. Du, N. Jillette, J. J. Zhu, S. Li, A. W. Cheng, CRISPR artificial splicing factors. Nat. Commun. 11, 2973 (2020).

22. E. J. Charles, S. E. Kim, G. J. Knott, D. Smock, J. Doudna, D. F. Savage, Engineering improved Cas13 effectors for targeted post-transcriptional regulation of gene expression. bioRxiv (2021), p. 2021.05.26.445687.

23. C. Wilson, P. J. Chen, Z. Miao, D. R. Liu, Programmable m6A modification of cellular RNAs with a Cas13-directed methyltransferase. Nat. Biotechnol. 38, 1431–1440 (2020).

24. K.-I. Miyazono, T. Miyakawa, Y. Sawano, K. Kubota, H.-J. Kang, A. Asano, Y. Miyauchi, M. Takahashi, Y. Zhi, Y. Fujita, T. Yoshida, K.-S. Kodaira, K. Yamaguchi-Shinozaki, M. Tanokura, Structural basis of abscisic acid signalling. Nature. 462, 609–614 (2009).

25. S. Kannan, H. Altae-Tran, X. Jin, V. J. Madigan, R. Oshiro, K. S. Makarova, E. V. Koonin, F. Zhang, Compact RNA editors with small Cas13 proteins. Nat. Biotechnol. 40, 194–197 (2022).

26. C. Xu, Y. Zhou, Q. Xiao, B. He, G. Geng, Z. Wang, B. Cao, X. Dong, W. Bai, Y. Wang, X. Wang, D. Zhou, T. Yuan, X. Huo, J. Lai, H. Yang, Programmable RNA editing with compact CRISPR– Cas13 systems from uncultivated microbes. Nat. Methods. 18, 499–506 (2021).

27. G. J. Knott, A. East-Seletsky, J. C. Cofsky, J. M. Holton, E. Charles, M. R. O’Connell, J. A. Doudna, Guide-bound structures of an RNA-targeting A-cleaving CRISPR-Cas13a enzyme. Nat. Struct. Mol. Biol. 24, 825–833 (2017).

28. C. Zhang, S. Konermann, N. J. Brideau, P. Lotfy, S. J. Novick, T. Strutzenberg, P. R. Griffin, P. D. Hsu, D. Lyumis, Structural basis for the RNA-guided ribonuclease activity of CRISPR-Cas13d. bioRxiv, 314401 (2018).

29. L. Toualbi, M. Toms, M. Moosajee, USH2A-retinopathy: From genetics to therapeutics. Exp. Eye Res. 201, 108330 (2020).

30. R. Koenekoop, M. Arriaga, K. M. Trzupek, J. Lentz, Usher Syndrome Type II (University of Washington, Seattle, 2023).

31. W. Li, X.-S. Jiang, D.-M. Han, J.-Y. Gao, Z.-T. Yang, L. Jiang, Q. Zhang, S.-H. Zhang, Y. Gao, J.-H. Wu, J.-K. Li, Genetic Characteristics and Variation Spectrum of USH2A-Related Retinitis Pigmentosa and Usher Syndrome. Front. Genet. 13, 900548 (2022).

32. T. S. Nambiar, L. Baudrier, P. Billon, A. Ciccia, CRISPR-based genome editing through the lens of DNA repair. Mol. Cell. 82, 348–388 (2022).

33. J. A. Hussmann, J. Ling, P. Ravisankar, J. Yan, A. Cirincione, A. Xu, D. Simpson, D. Yang, A. Bothmer, C. Cotta-Ramusino, J. S. Weissman, B. Adamson, Mapping the genetic landscape of DNA double-strand break repair. Cell. 184, 5653–5669.e25 (2021).

34. D. S. Mackay, A. D. Borman, R. Sui, L. I. van den Born, E. L. Berson, L. A. Ocaka, A. E. Davidson, J. R. Heckenlively, K. Branham, H. Ren, I. Lopez, M. Maria, M. Azam, A. Henkes, E. Blokland, R. Qamar, A. R. Webster, F. P. M. Cremers, A. T. Moore, R. K. Koenekoop, [LCA5 Study Group (see acknowledgements for Universities), S. Andreasson, E. de Baere, J. Bennett, G. J. Chader, W. Berger, I. Golovleva, J. Greenberg, A. I. den Hollander, C. C. W. Klaver, B. J. Klevering, B. Lorenz, M. N. Preising, R. Ramsear, L. Roberts, R. Roepman, K. Rohrschneider, B. Wissinger, Screening of a large cohort of leber congenital amaurosis and retinitis pigmentosa patients identifies novel LCA5 mutations and new genotype-phenotype correlations. Hum. Mutat. 34, 1537–1546 (2013).

35. A. B. Popejoy, S. M. Fullerton, Genomics is failing on diversity. Nature Publishing Group UK (2016),, doi:10.1038/538161a.

36. S. Picelli, O. R. Faridani, A. K. Björklund, G. Winberg, S. Sagasser, R. Sandberg, Full-length RNA-seq from single cells using Smart-seq2. Nat. Protoc. 9, 171–181 (2014).

37. S. Nurk, D. Meleshko, A. Korobeynikov, P. A. Pevzner, metaSPAdes: a new versatile metagenomic assembler. Genome Res. 27, 824–834 (2017).

38. D. Hyatt, G.-L. Chen, P. F. Locascio, M. L. Land, F. W. Larimer, L. J. Hauser, Prodigal: prokaryotic gene recognition and translation initiation site identification. BMC Bioinformatics. 11, 119 (2010).

39. K. Katoh, D. M. Standley, MAFFT multiple sequence alignment software version 7: improvements in performance and usability. Mol. Biol. Evol. 30, 772–780 (2013).

40. B. Q. Minh, H. A. Schmidt, O. Chernomor, D. Schrempf, M. D. Woodhams, A. von Haeseler, R. Lanfear, IQ-TREE 2: New Models and Efficient Methods for Phylogenetic Inference in the Genomic Era. Mol. Biol. Evol. 37, 1530–1534 (2020).

41. D. D. Kang, F. Li, E. Kirton, A. Thomas, R. Egan, H. An, Z. Wang, MetaBAT 2: an adaptive binning algorithm for robust and efficient genome reconstruction from metagenome assemblies. PeerJ. 7, e7359 (2019).

42. Y.-W. Wu, B. A. Simmons, S. W. Singer, MaxBin 2.0: an automated binning algorithm to recover genomes from multiple metagenomic datasets. Bioinformatics. 32, 605–607 (2016).

43. J. Alneberg, B. S. Bjarnason, I. de Bruijn, M. Schirmer, J. Quick, U. Z. Ijaz, N. J. Loman, A. F. Andersson, C. Quince, CONCOCT: Clustering cONtigs on COverage and ComposiTion. arXiv [q-bio.GN] (2013), (available at http://arxiv.org/abs/1312.4038).

44. D. H. Parks, M. Imelfort, C. T. Skennerton, P. Hugenholtz, G. W. Tyson, CheckM: assessing the quality of microbial genomes recovered from isolates, single cells, and metagenomes. Genome Res. 25, 1043–1055 (2015).

45. C. Sieber, Dereplication, Aggregation and Scoring Tool (DAS Tool) v1.0 (Lawrence Berkeley National Laboratory (LBNL), Berkeley, CA (United States), 2017; https://www.osti.gov/biblio/1345357).

