## Supplement for "Programmable multi-kilobase RNA editing using CRISPR-mediated trans-splicing"

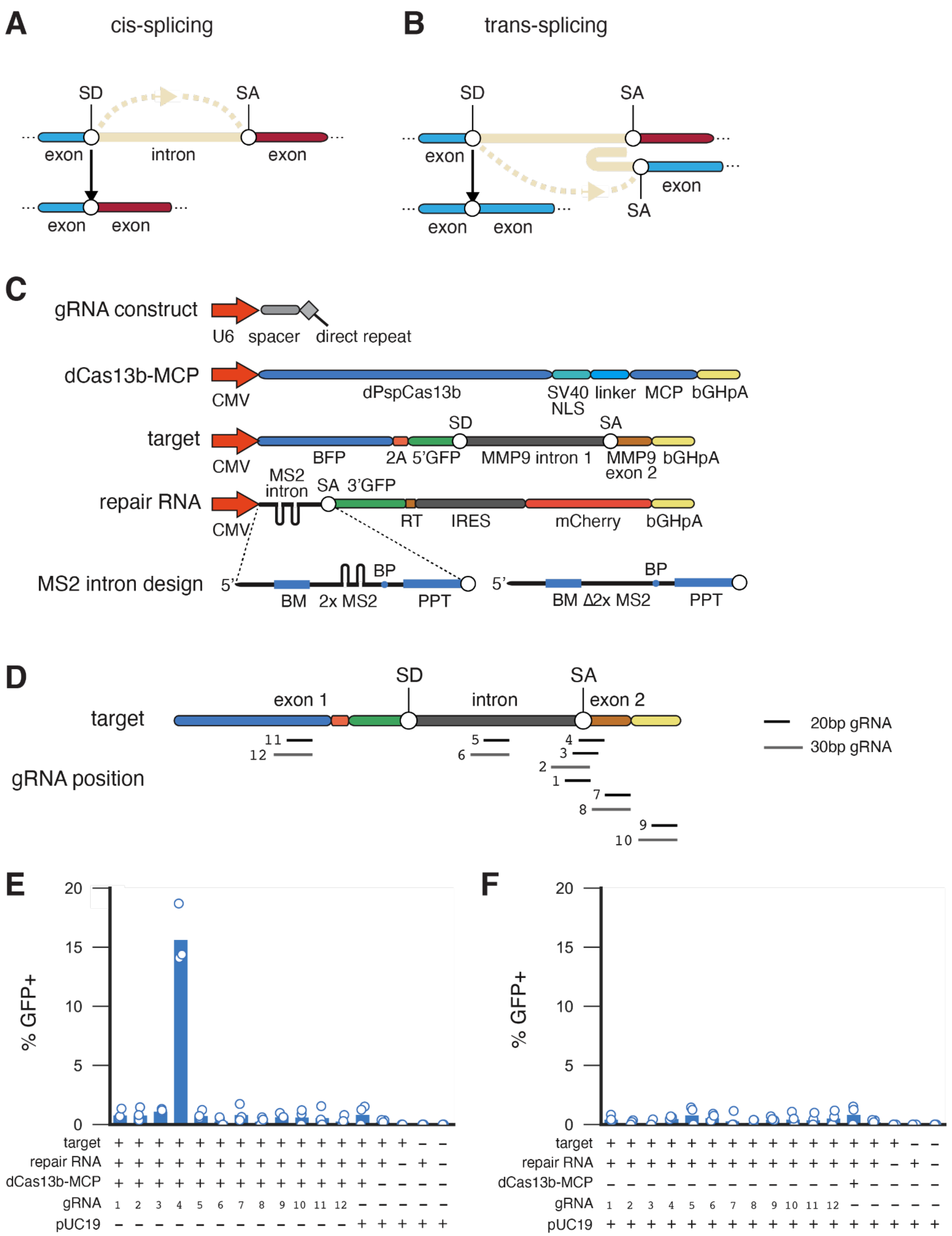
A
cis-splicing
SD
SA
exon
intron
exon
exon
exon
B
trans-splicing
SD
SA
exon
exon
SA
exon
exon
C
gRNA construct
U6
spacer
direct repeat
dCas13b-MCP
CMV
dPspCas13b
SV40 NLS
linker
MCP
bGHpA
target
CMV
BFP
2A
5'GFP
SD
MMP9 intron 1
SA
MMP9 exon 2
bGHpA
repair RNA
CMV
MS2 intron
SA
3'GFP
RT
IRES
mCherry
bGHpA
MS2 intron design
5'
BM
2x MS2
BP
PPT
5'
BM
Δ2x MS2
BP
PPT
D
target
exon 1
SD
intron
SA
exon 2
gRNA position
20bp gRNA
30bp gRNA
E
% GFP+
target
repair RNA
dCas13b-MCP
gRNA
pUC19
F
% GFP+
target
repair RNA
dCas13b-MCP
gRNA
pUC19

**Figure S1.** (**A**) RNA cis-splicing. (**B**) RNA trans-splicing. (**C**) DNA constructs for trans-splicing reporter and Splice Editor systems. The synthetic MS2 intron in the repRNA contains a binding motif (BM) that hybridizes to the target pre-mRNA, along with a branch point (BP) and polypyrimidine tract (PPT). (**D**) Schematic of gRNA target sites along target construct. gRNAs 1-4 targeted the splice acceptor (SA) site, while gRNAs 5 and 6 targeted the intron. gRNAs 7 and 8 targeted MMP9 exon 2, while gRNAs 9 and 10 targeted bGHpA. gRNAs 11 and 12 targeted BFP in the first exon. (**E**) Activity of targeted RNA trans-splicing with dCas13b-MCP Splice Editor in mammalian HEK293FT cells (n=3 biological replicates). (**F**) Activity of RNA trans-splicing without dCas13b-MCP in mammalian HEK293FT cells (n=3 biological replicates), demonstrating activity of the repRNA alone.

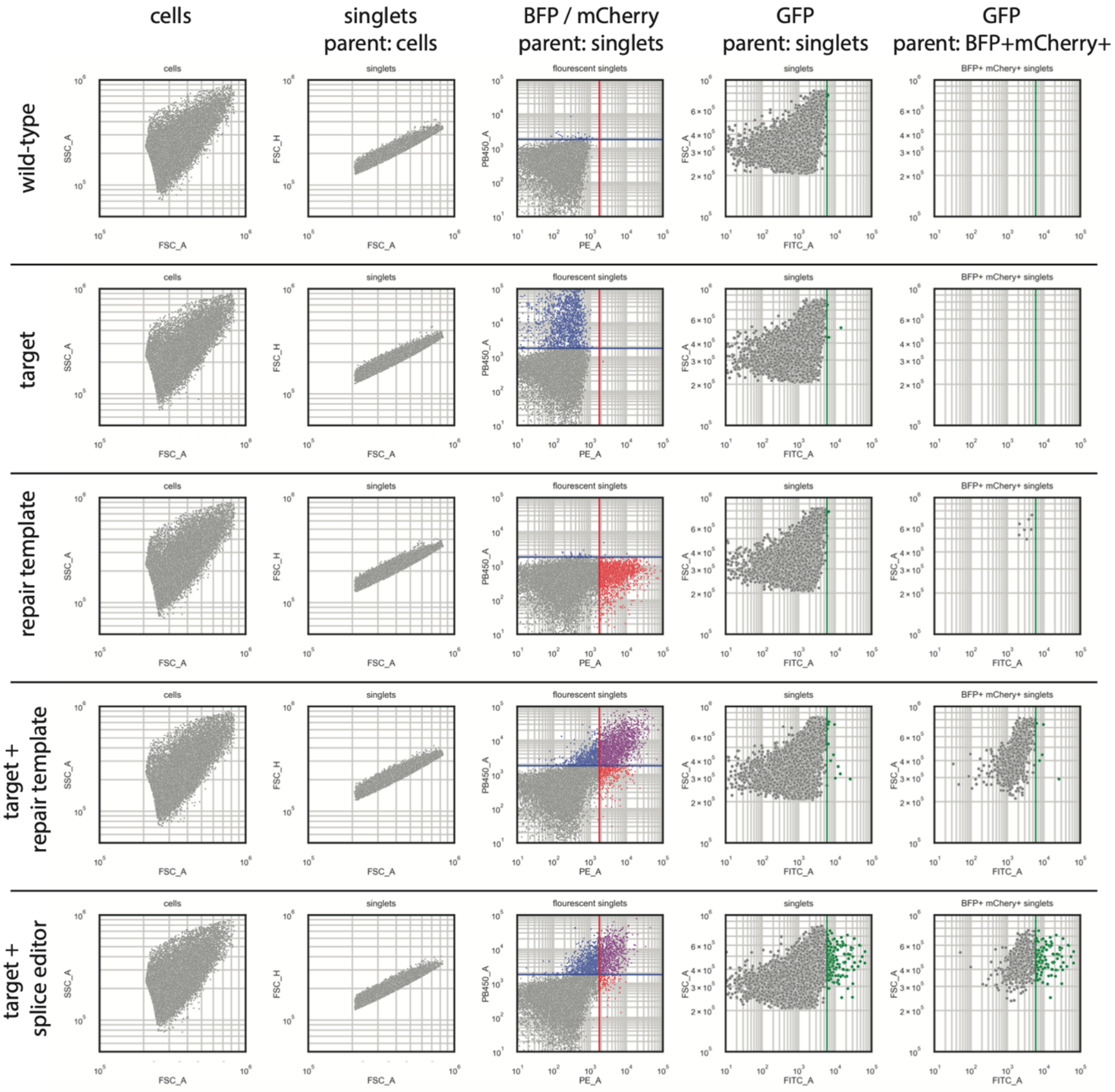


**Figure S2. Flow cytometry gating strategy.** Cells were gated from side scatter (area) and forward scatter (area), and singlets were gated on forward scatter (height vs. area). BFP and mCherry signal was detected relative to background and then BFP+mCherry+ singlets were considered for GFP analysis, which was gated on GFP fluorescence above the wild-type singlets.

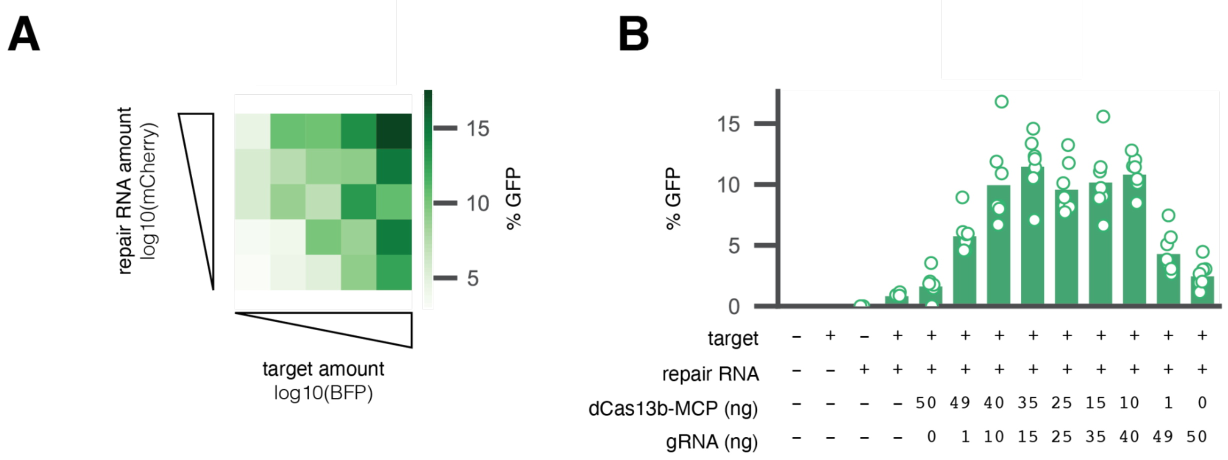


**Figure S3. 3' splice editor component titration.**
(**A**) Trans-splicing activity is dependent on target and repRNA amounts, as measured by flow cytometry. Increased repRNA amounts led to higher editing efficiencies. (**B**) Splice editing showed similar activities across a range of gRNA and dCas13b-MCP amounts in transfection experiments, indicating that activity is not gRNA or dCas13b-MCP limited.

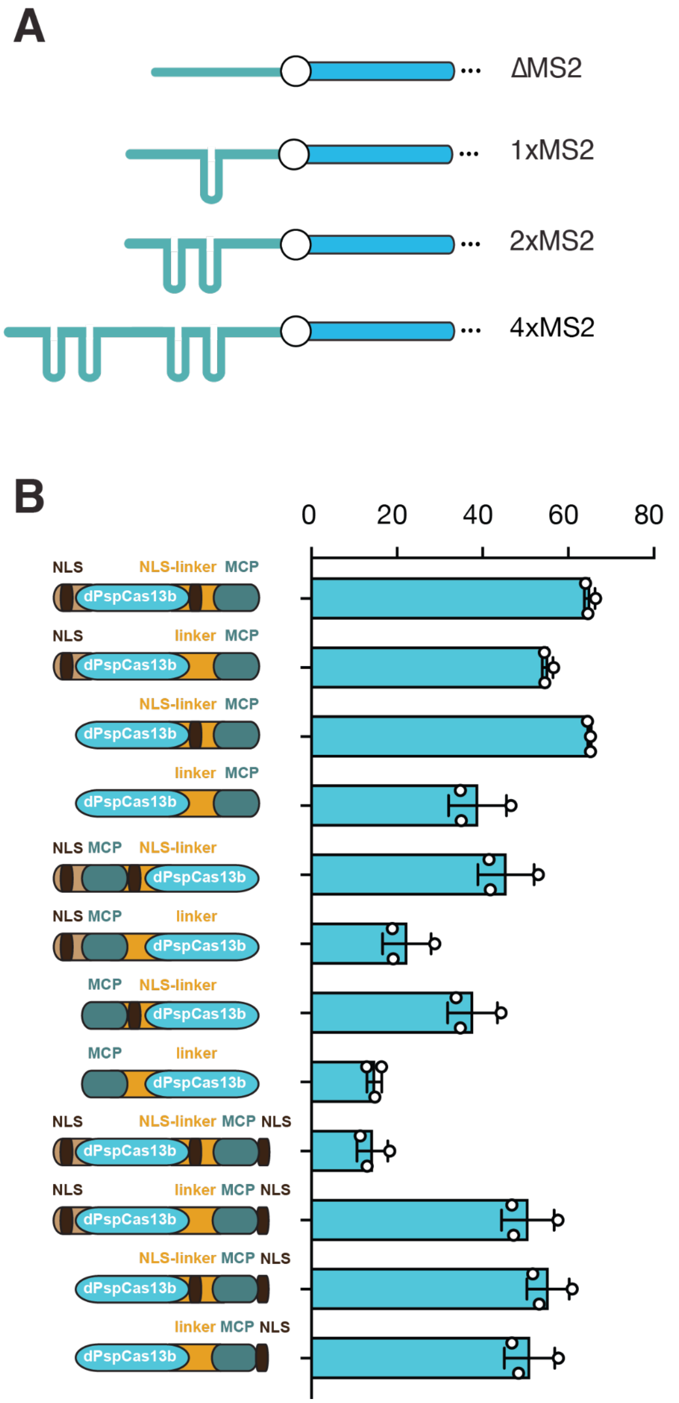


**Figure S4. (A)** Schematic of different repRNA hairpin compositions tested for their impact on Splice Editing activity. Increasing the number of hairpins led to improved Splice Editing activity as observed in Figure 1H. **(B)** Different Splice Editor architectures were generated via a combinatorial optimization library consisting of different RBP fusions and NLS positions.

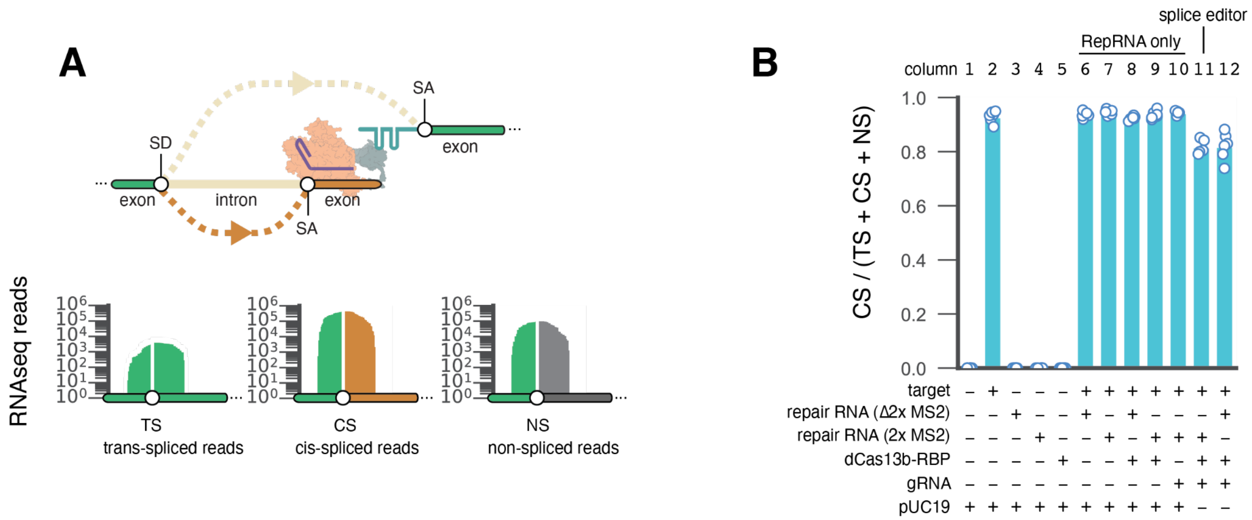


**Figure S5. RNA-seq validation of trans-splicing and cis-splicing.**
(**A**) Full-length RNA-seq was conducted on 293FT cells transfected with dCas13b-MCP, gRNA, and the split GFP reporter system (target, repRNA w/ 2x MS2). Reads spanning all possible junctions were filtered and subsequently mapped onto all possible junction sequences, namely, trans-spliced junction (TS), cis-spliced junction (CS) and non-spliced junction (NS). The mapping positions were then plotted to illustrate reads spanning possible junctions. (**B**) The fraction of spliced reads that were cis-spliced was calculated and plotted for all transfection conditions. Results show that dCas13 inhibits cis-splicing in the presence of a targeting gRNA.

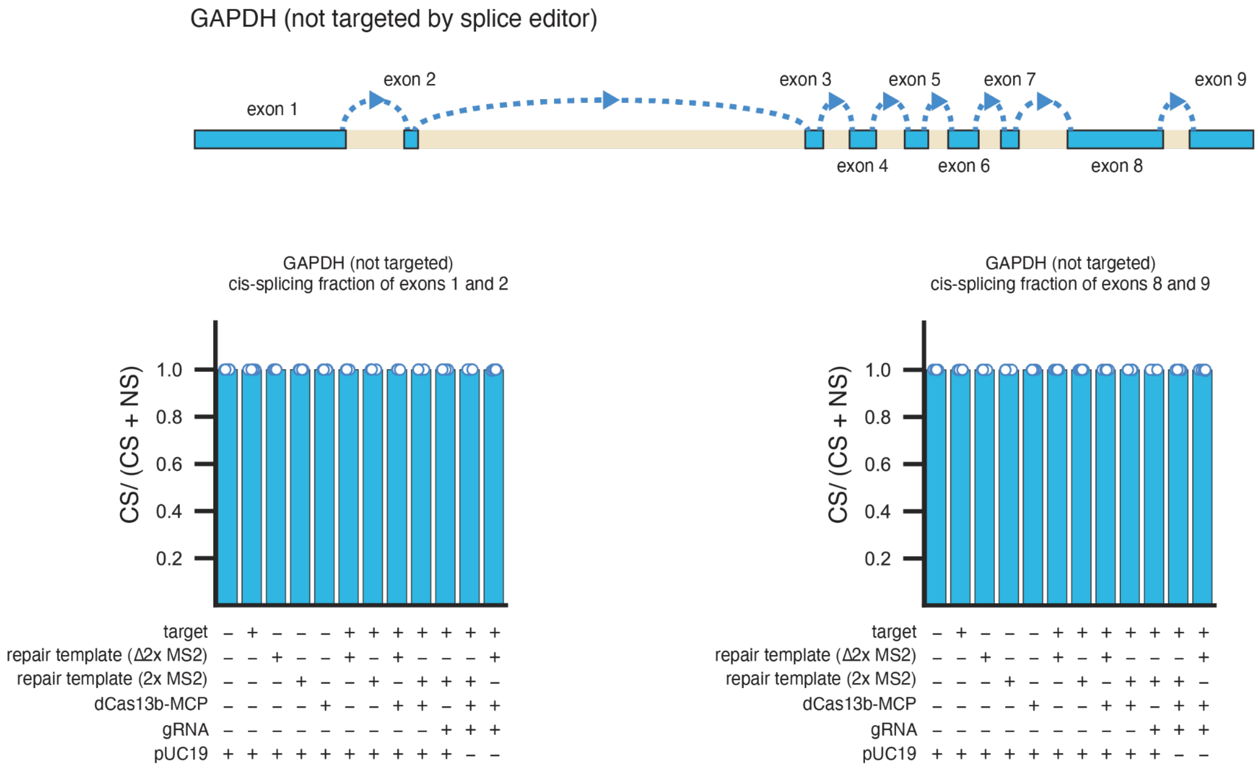


**Figure S6. Cis-splicing rates of non-targeted GAPDH pre-mRNA.**
Full-length RNA-seq was conducted on 293FT cells transfected with dCas13b-MCP, gRNA, and the split GFP reporter system (target, repRNA w/ 2x MS2). Reads spanning cis-spliced (CS) and non-spliced (NS) junctions were filtered and counted to measure cis-splicing rates for exons 1 and 2, as well as exons 8 and 9. Results demonstrate that non-targeted GAPDH pre-mRNA cis-splicing rates were not altered in the presence of Splice Editing components.

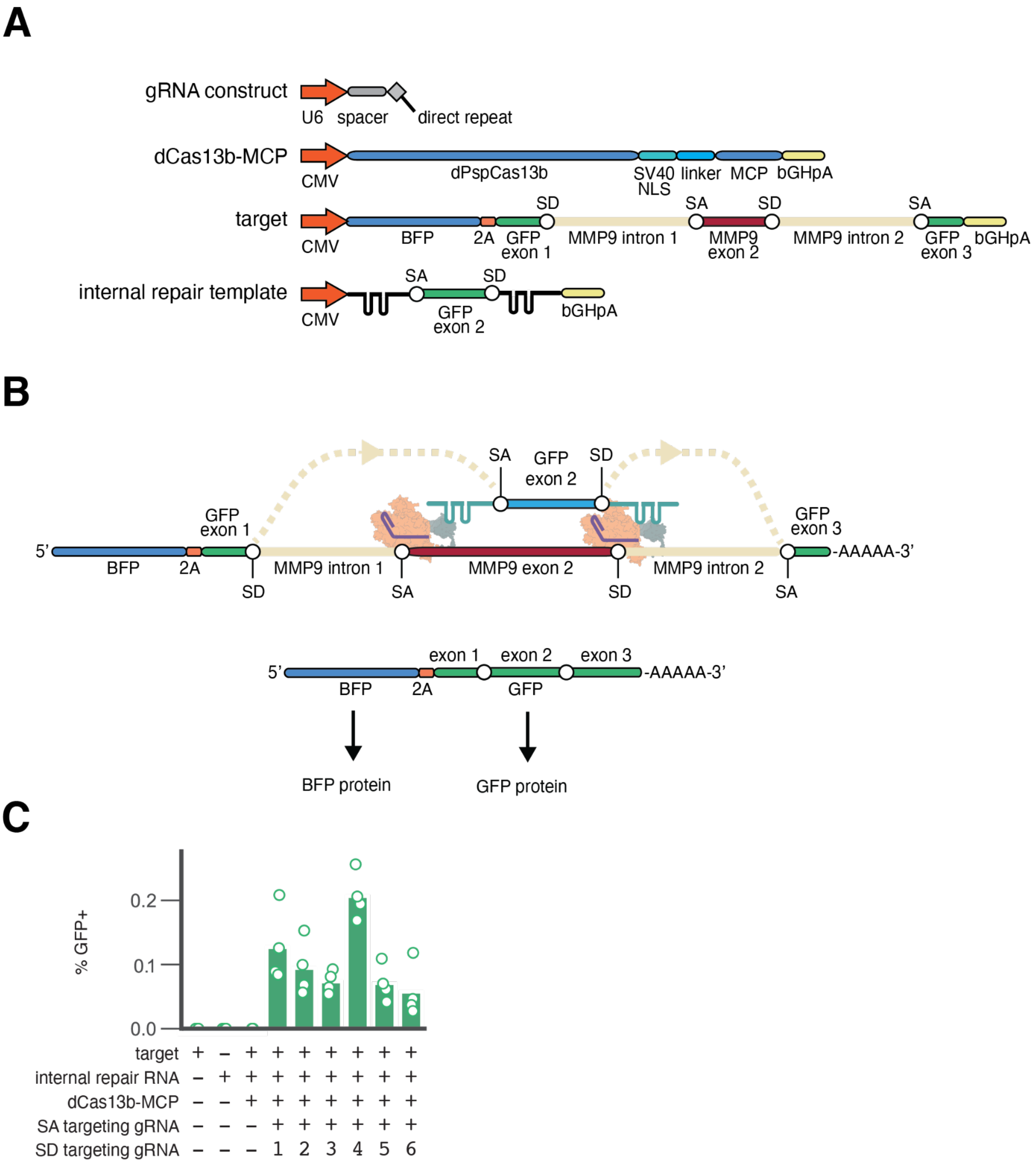


**Figure S7. Internal exon repair with splice editors.**
(**A**) Standard gRNA constructs were used, along with CMV-dPspCas13b-MCP, and an internal GFP repair reporter to test whether internal exon repair is possible with CRISPR-mediated trans-splicing. (**B**) A double trans-splicing event mediated by a splice editor is measured through the detection of GFP. (**C**) Internal exon repair was only detected in cases where there was a splice editor and a splice donor (SD) targeting gRNA and a splice acceptor (SA) targeting gRNA

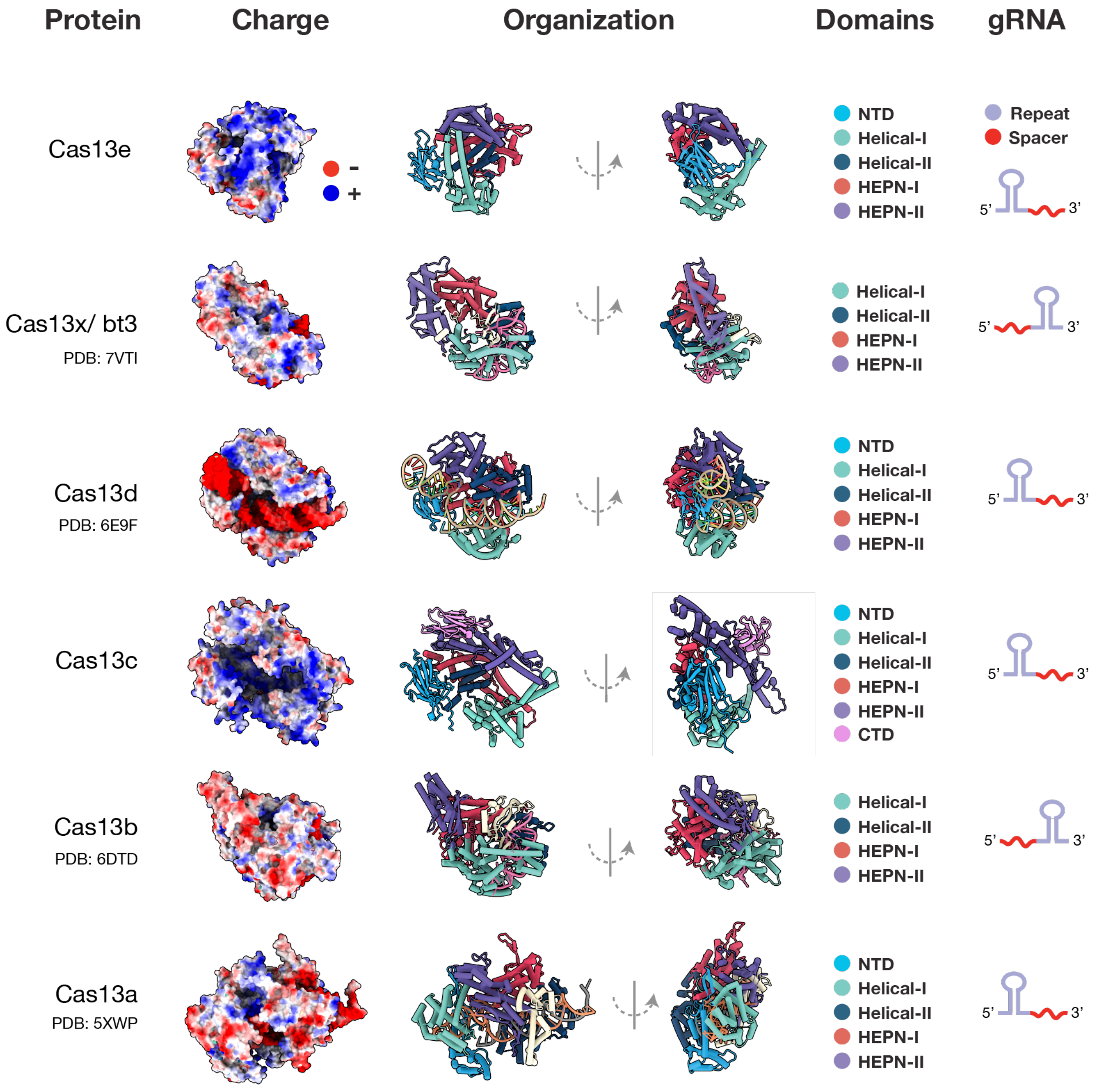


**Figure S8.** The architecture of Cas13e as compared to other type VI (Cas13) proteins. Available ternary or binary structures of type VI effectors in the crRNA, or crRNA/ target RNA bound states are shown. The charge across each structure is shown, with negative charges in red and positive charges in blue. Domains are color-coded according to the legends shown and two displays of the domain organization shown. The gRNA orientation for each enzyme is shown, with CRISPR repeats colored in purple and spacers in red. The following models were used to prepare the figure: Cas13e and Cas13c predicted structures (this study); Cas13a (PDB-ID: 5XWP); Cas13b (PDB-ID: 6DTD); Cas13d (PDB-ID: 6E9F); Cas13bt3 (PDB-ID: 7VTI).
